# Nucleolin-RNA interaction modulates rotavirus replication

**DOI:** 10.1101/2023.10.02.560526

**Authors:** Jey Hernández-Guzmán, Carlos F. Arias, Susana López, Carlos Sandoval-Jaime

## Abstract

Rotavirus infection is a leading cause of gastroenteritis in children worldwide, the genome of this virus is composed of eleven segments of dsRNA packed in a triple-layered protein capsid. Here, we investigated the role of nucleolin, a protein with diverse RNA-binding domains, on rotavirus infection. Knocking down the expression of nucleolin in MA104 cells by RNA interference resulted in a remarkable 5-fold increase in the production of infectious rhesus rotavirus (RRV) progeny, accompanied by an elevated synthesis of viral mRNA and genome copies. Further analysis unveiled an interaction between rotavirus segment 10 (S10) and nucleolin, potentially mediated by G-quadruplex domains on the viral genome. To determine whether the nucleolin-RNA interaction regulates RRV replication, MA104 cells were transfected with AGRO100, a compound that forms G4 structures and selectively inhibits nucleolin-RNA interactions by blocking the RNA binding domains. Under these conditions, viral production increased by 1.54-fold, indicating the inhibitory role of nucleolin on the yield of infectious viral particles. Furthermore, G4 sequences were identified in all eleven RRV dsRNA segments, and transfection of oligonucleotides representing G4 sequences in RRV S10 induced a significant increase in viral production. These findings show that rotavirus replication is negatively regulated by nucleolin through direct interaction with the viral RNAs by sequences forming G4 structures.

**Importance:** Viruses rely on cellular proteins to carry on their replicative cycle. In the case of rotavirus, the involvement of cellular RNA-binding proteins during the replicative cycle is a poorly studied field. In this work, we demonstrate for the first time the interaction between nucleolin and viral RNA of rotavirus RRV. Nucleolin is a cellular protein that has a role in the metabolism of ribosomal rRNA, and ribosome biogenesis, which seems to have regulatory effects on the quantity of viral particles and viral RNA copies of rotavirus RRV. Our study adds a new component to the current model of rotavirus replication, where cellular proteins can have a negative regulation on rotavirus replication.

## Introduction

Rotavirus-induced gastroenteritis poses a significant global health burden, particularly affecting children under five years old. Although the availability of recombinant vaccines, such as Rotateq and Rotarix, has contributed to a reduction in the incidence of severe cases of diarrhea, this virus continues to cause substantial morbidity and mortality worldwide (Burnett et al., 2018). Rotaviruses are members of the Reoviridae family, non-enveloped viruses characterized by a triple-layered protein capsid. Their genome is composed by 11 segments of double-stranded RNA (dsRNA) that encode six structural viral proteins (VP1, VP2, VP3, VP4, VP6, and VP7) and six nonstructural proteins (NSP1, NSP2, NSP3, NSP4, NSP5, and NSP6) (Crawford et al., 2017).

During infection, rotavirus employs intricate strategies to translate viral proteins and replicate its genome by modulating the cellular environment and hijacking host proteins (Crawford et al., 2017). While substantial knowledge exists regarding the functions of viral proteins, our understanding of the cellular proteins involved in the rotavirus replication cycle remains limited. Several proteins with RNA binding domains have been found to change their cellular localization upon rotavirus infection, silencing the expression of HUR, hnRNP L, I, D, ATP5B and Ago2 increases the production of viral progeny (Dhillon et al., 2018; Oceguera et al., 2018; Ren et al., 2019), suggesting that they have a role during the replication cycle of rotavirus.

Nucleolin, a eukaryotic, mostly nuclear phosphoprotein, possesses a central region with an RNA-binding domain (RBD) comprising four RNA recognition motifs that enable interactions with RNA molecules (Tajrishi et al., 2011). This protein plays diverse roles in ribosome assembly, rDNA transcription, and the modification and processing of pre-rRNA (Tajrishi et al., 2011). Nucleolin has been reported to participate in cell binding and entry of several RNA viruses, thereby influencing their initial infection process (Bose et al., 2004; Tayyari et al., 2011; Zhu et al., 2018). In addition, it has been implicated in viral replication and translation of caliciviruses, (Cancio-Lonches et al., 2011; Hernández et al., 2016), and the assembly of dengue virus particles (Balinsky et al., 2013).

In this work, we report that the number or viral RNA copies and the viral progeny increases when the expression of nucleolin was knocked down in MA104 cells. Immunoprecipitation assays reveal that nucleolin interacts with the RNA of rotavirus more likely through its binding to G-quadruplex RNA structures. Accordingly, using synthetic sequences predicted to form G-quadruplex structures and that mimic sequences in the rotavirus gene segment 10, we were able to confirm that these regions block the binding of nucleolin causing an increase in the production of infectious viral particles.

## Materials and methods

### Cell Culture and Viruses

The African green monkey kidney epithelial cell line MA104 (ATCC) was cultivated in Dulbecco’s modified Eagle’s medium (DMEM) (Thermo Scientific HyClon, Logan, UT) and supplemented 5% fetal bovine serum (FBS) (Biowest, Kansas, MO) at 37°C in a 5% CO_2_ atmosphere. The rhesus rotavirus strain RRV utilized in this investigation was initially donated by H. B. Greenberg (Stanford University, Stanford, CA). The virus was propagated in MA104 cells, and the resultant virus was titrated and stored at -70°C.

### Antibodies

Polyclonal antibodies targeting purified rotavirus RRV triple-layered particles (TLP’s), and vimentin were generated in rabbits, following previously described protocols (Lopez et al., 2005). Rabbit polyclonal sera against NSP3 was prepared as described previously (Montero et al., 2006). Monoclonal antibodies specific to nucleolin and PPI were obtained from Santa Cruz Biotechnology (Santa Cruz, CA). Horseradish peroxidase-conjugated goat anti-rabbit polyclonal antibody was sourced from KPL (Gaithersburg, MD), while horseradish peroxidase-conjugated goat anti-mouse antibody was acquired from Millipore Merck KGaA (Darmstadt, Germany). Streptavidin Magnetic Beads were procured from New England Biolabs (Massachusetts, US).

### Cell Infection and virus titration

Prior to infection, the RRV was activated by incubating with trypsin (10 μg/mL, Gibco, Life Technologies, Carlsbad, CA) for 30 minutes at 37°C. Monolayers of MA104 cells were grown in 48-well plates and infected with a multiplicity of infection (MOI) of 3 for 15 hours. The infected cells were lysed by three freeze-thaw cycles. The viral lysate’s infection titers were determined using an immunoperoxidase assay. Briefly, confluent cells in 96-well plates were incubated with two-fold serial dilutions of the viral lysate (previously activated with trypsin as above) for 60 minutes at 37°C. After the adsorption period, the virus inoculum was removed, the cells were washed once, and fresh MEM was added. The infection was allowed to proceed for 15 hours at 37°C. RRV-infected cells were detected using a rabbit hyperimmune serum to rotavirus in an immunoperoxidase focus detection assay, following previously described protocols (Gutierrez et al., 2010).

### siRNA transfection

Small interfering RNAs (siRNAs) targeting nucleolin (SMARTpool siGENOME Human Ncl, M-003854-01-0005) and a “NonTargeting” (SMARTpool siGENOME NonTargeting, D-001210-04-20) control siRNAs were obtained from GE Healthcare Dharmacon (Lafayette, CO). The siRNA transfection was performed in MA104 cells grown in 48-well plates using a reverse transfection method, as previously described (Gutierrez et al., 2010). Briefly, the transfection mix [15 µL Oligofectamine/mL (Invitrogen, Eugene, OR), mixed with 2.5 μM of the appropriate siRNA] was added to the well, and 20,000 cells/well were subsequently added and incubated for 16 hours at 37°, the transfection mix was then replaced by MEM, and the cells were further incubated for up to 72 hours at 37°C.

### Immunoblot Analysis

For immunoblot analysis, cells were lysed using Laemmli sample buffer (50 mM Tris-HCl pH 6.8, 2% SDS, 0.1% bromophenol blue, 10% glycerol, 5% β-mercaptoetanol) and denatured by boiling at 95°C for 5 minutes. The protein lysates were subjected to electrophoresis on 10% SDS-polyacrylamide gels, followed by transfer of proteins to an Immobilon membrane (Millipore) using CAPS buffer (CAPS 10mM, 10% methanol). The membrane was then blocked for 1 hour at room temperature with 5% skim milk in PBS-0.1% Tween. Next, the membrane was incubated with primary antibodies diluted 1:3000 in a solution of 1% skim milk in PBS-0.1% Tween, for 1 hour at room temperature. The binding of the primary antibodies was detected using horseradish peroxidase-conjugated anti-rabbit immunoglobulins (Thermo Scientific, Waltham, MA) at a 1:3000 dilution, or with peroxidase conjugated anti-mouse immunoglobulins (Thermo Scientific, Waltham, MA,) at a 1:3000 dilution. Protein bands were developed using the Western Lightning system (PerkinElmer, MA). Densitometry analysis of the Western blot images was performed using Image J software (Abràmoff et al., 2004).

### Real-time RT-PCR

Confluent MA104 cells were seeded in 48-well plates and infected with RRV at a multiplicity of infection (MOI) of 3. The cells were lysate at different times post-infection using Trizol reagent, and the RNA was extracted using the manufacturer instructions. The primers used for the amplification of gene 10 of rotavirus were previously described as oligos forward: TCCTGGAATGGCGTATTTTC and reverse: GAGCAATCTTCATGGTTGGAA (Ayala-Breton et al., 2009). Reverse transcription-polymerase chain reaction (RT-PCR) was employed to specifically determine the levels of each RNA strand, following the previously described method (Ayala-Breton et al., 2009). The quantitative analysis of data was performed using the ABI Prism 7000 analysis software program. After PCR amplification, the fluorescence values of all samples between the logarithmic phases of the amplification curves were used to set a cutoff line using the ABI Prism software. The logarithm of the concentration of each sample was plotted against the cycle number, where the amplification curve of the sample reached the cutoff line (CT). The amount of positive or negative strand RNA from experimental samples was determined by extrapolating the CT value into the corresponding standard curve. The standard curve was prepared using a plasmid of DNA with the NSP4 gene sequence of rotavirus RRV present at 10^1^-10^11^ copies number. Experimental samples were collected at 0, 4, 8, and 12 hpi.

### Immunoprecipitation Assays

Confluent MA104 cells were grown in 6-well plates and infected with RRV at an MOI of 5. At 8 hpi, the cells were washed with PBS-EDTA (0.2 mM) and harvested by scraping. The resulting cell pellet was centrifuged for 10 minutes at 1000 RPM at 4°C and subsequently washed twice with 2 mL of PBS at 4°C, followed by centrifugation for 10 minutes at 1000 RPM. The resulting pellet was then resuspended in 1 mL of RIPA buffer (50 mM Tris-HCl at pH 7.5, 1% Triton, 0.5% sodium deoxycholate, 0.005% SDS, 1 mM EDTA, and 150 mM NaCl), containing proteases inhibitors (Roche, Basilea) and RNases inhibitors. (NEB, MA) and subjected to three rounds of sonication (20 seconds each). The resulting cell lysate was centrifuged for 10 minutes at 14,000 RPM, and the supernatants were subsequently clarified with 50 µL of magnetic beads coupled to protein G (Dynabeads Protein G-LS10003D, Thermo Scientific) at 4°C for 1 h with agitation. 15 µL of magnetic beads Dynabeads were pre-treated with 10 mg/mL of tRNA, and antibodies were added to the pre-cleared lysate and incubated overnight at 4°C with agitation. After this time, the beads were washed four times with RIPA buffer. The beads were split in two separate tubes: the first tube was used for RNA extraction using Trizol, while the second tube was used to identify the immunoprecipitated proteins through western blot.

### Analysis of Putative G-Quadruplex (G4) Domains

The QGRS mapper web server (Kikin et al., 2006) was utilized to search for putative G-quadruplex domains. The positive polarity RNA strand sequences of the eleven segments of rotavirus RRV were examined. The following parameters were employed: maximum domain length of 45 nucleotides, mini groups of guanines together of 2, and possible loop sizes of up to 36 nucleotides (Kikin et al., 2006). The search was performed using the QGRS mapper web server, available at https://bioinformatics.ramapo.edu/QGRS/analyze.php.

### G-quadruplex formation and transfection

The G-quadruplex-forming sequences used in these assays were:

AGRO100: 5’-GGTGGTGGTGGTTGTGGTGGTGGTGG-3’
CRO26: 5’-CCTCCTCCTCCTTCTCCTCCTCCTCC-3’
G4-1: 5’-GGAGGATCCTGGAATGG-3’
G4-2: 5’-GGTTGAGCTGCCGTCGTCTGTCTGCGGAAGCGGCGG-3’
G4-3: 5’-GGACGTTAATGGAAGGAACGG-3’

The G-quadruplex folding was carried out as described previously (Hacht *et. al.*, 2014). Briefly, each G4 oligonucleotide was suspended in folding buffer (10mMTris-HCl, pH 7.5, 100mMKCl and 0.1 mM EDTA), and the formation of secondary structures was achieved by heating the ssDNA to 95◦C for 5 min and cooling it down to 4◦C in steps of 2◦C per minute. The folded G4 sequences were transfected into MA104 cells in 48-well plates as described previously for the siRNA transfections.

### Statistical Analysis

The statistical significance of the experiments was evaluated using the Mann-Whitney U test with a significance level of 95%. The GraphPad Prism 5 program was employed for the statistical analysis. The asterisks displayed in the graphs represent different degrees of statistical significance: *=p<0.05, **=p<0.01, and ***=p<0.001.

## Results

### Nucleolin negatively regulates rotavirus replication

To advance in the characterization of the role of RNA-binding cellular proteins on rotavirus replication, we evaluated the role of nucleolin, an RNA binding protein known to interact with several RNA viruses, on the replication cycle of rotaviruses. For this, the effect of silencing the expression of nucleolin by RNAi on the yield of infectious rotavirus progeny during a single replicative cycle was evaluated. In these assays, MA104 cells seeded in 48-well plates were transfected with a pool of siRNAs specifically targeting nucleolin and 72 h post-transfection, the cells were infected with rhesus rotavirus (RRV) at a MOI of 3. At 15 h post-infection (hpi), the infected cells were harvested, and the viral titer of the virus produced under these conditions was determined by an immunoperoxidase assay (Gutierrez et al., 2010) (Fig. 1A). We found a 5-fold increase in the viral titer obtained from cells in which nucleolin was silenced as compared to cells transfected with a control, non-targeting (NT) siRNA. This observation suggests that nucleolin is a negative regulation factor to produce rotavirus infectious particles (Fig. 1A).

**Figure 1.**
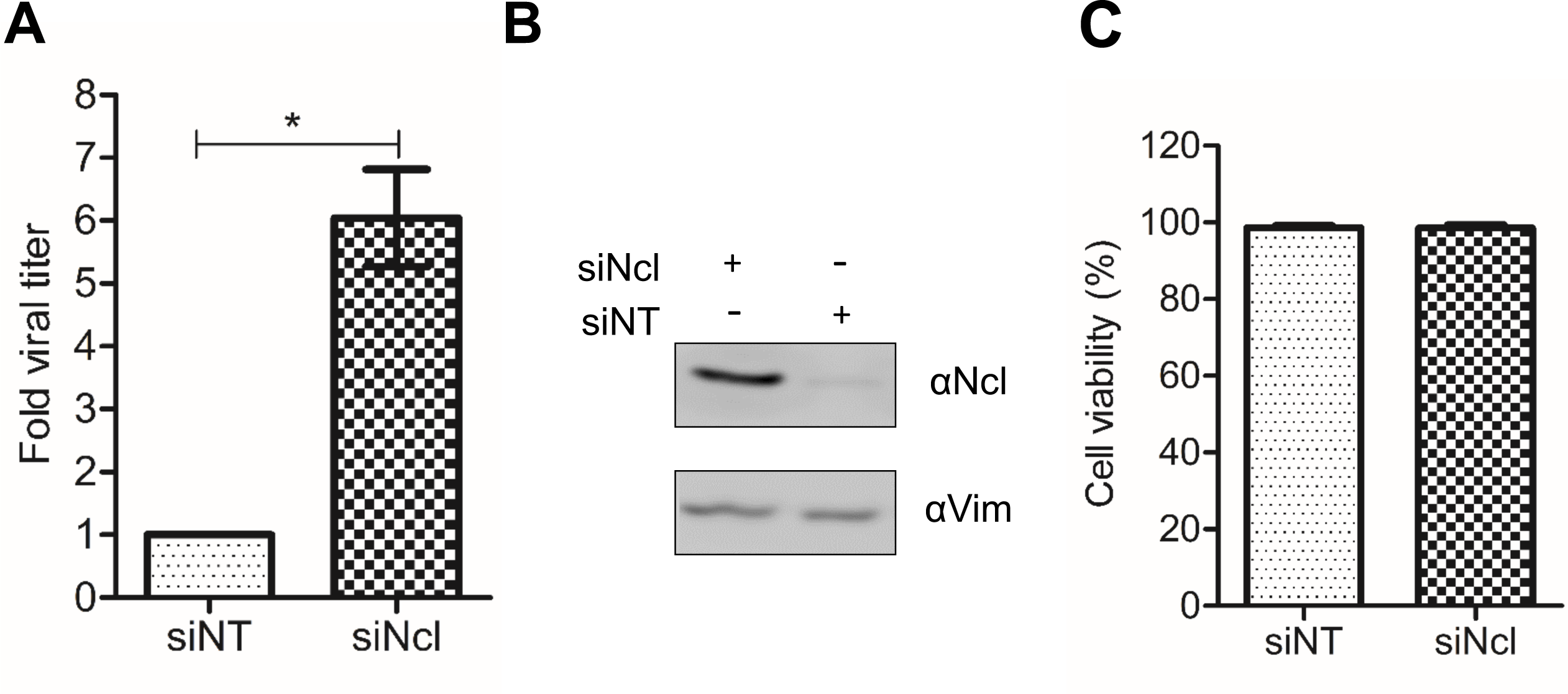
Knockdown of nucleolin expression promotes an increase in rotavirus titers. A) MA104 cells grown in 48 wells were transfected with either non-targeting siRNA (NT) or nucleolin siRNA (Ncl). At 72 hours post-transfection (hpt), cells were infected with RRV at an MOI of 3, harvested at 15 hpi, and subjected to three cycles of freezing and thawing. The virus titer in the cleared cell lysates was determined by an immunoperoxidase assay, as described under Materials and Methods. Data is expressed as fold increases in which the titer obtained from the cells transfected with the NT siRNA was used to normalize. Data shown represent the arithmetic mean ± standard deviation of three independent experiments. B) Cells transfected with the indicated siRNAs were harvested at 72 hpt, and the proteins were resolved by SDS-10% PAGE, transferred to a nitrocellulose membrane, and detected by immunoblot analysis, using antibodies to nucleolin (Ncl) and Vimentin (Vim) as a loading control. Densitometry analysis of the Ncl and Vim bands in the Western blot was performed using ImageJ software. C) Cell viability was assessed using the LDH release assay following the manufacturer’s specifications. The arithmetic means ± standard deviation of three independent experiments is shown.

The efficacy of the knockdown of nucleolin was found to be 91%, as evaluated by densitometry of western blots developed with an anti-nucleolin antibody, normalized to the loading control, using an anti-vimentin antibody (vimentin) (Fig. 1B). Under the conditions studied, no discernible changes in the cell viability, as determined by LDH release assay, were detected in cells treated with the siRNA directed to nucleolin (Figure 1C).

### Nucleolin has no influence on rotavirus cell entry or viral protein synthesis

To define the step of the virus replication cycle affected by the knockdown of nucleolin, we evaluated the initial stages of virus replication. For this, MA104 cells that had been previously treated with either nucleolin or NT siRNAs were infected with RRV at an MOI of 3. At 15 hpi, the cells were fixed and stained with a rabbit polyclonal antibody TLP’s raised against purified rotavirus particles, and the number of infected cells was quantified by an immunoperoxidase assay (see Material and Methods). The viral titer measured in both, nucleolin-silenced and NT control cells, exhibited no statistically significant difference between the two groups (Fig. 2A), indicating that reduction in nucleolin does not impact virus cell entry.

**Figure 2.**
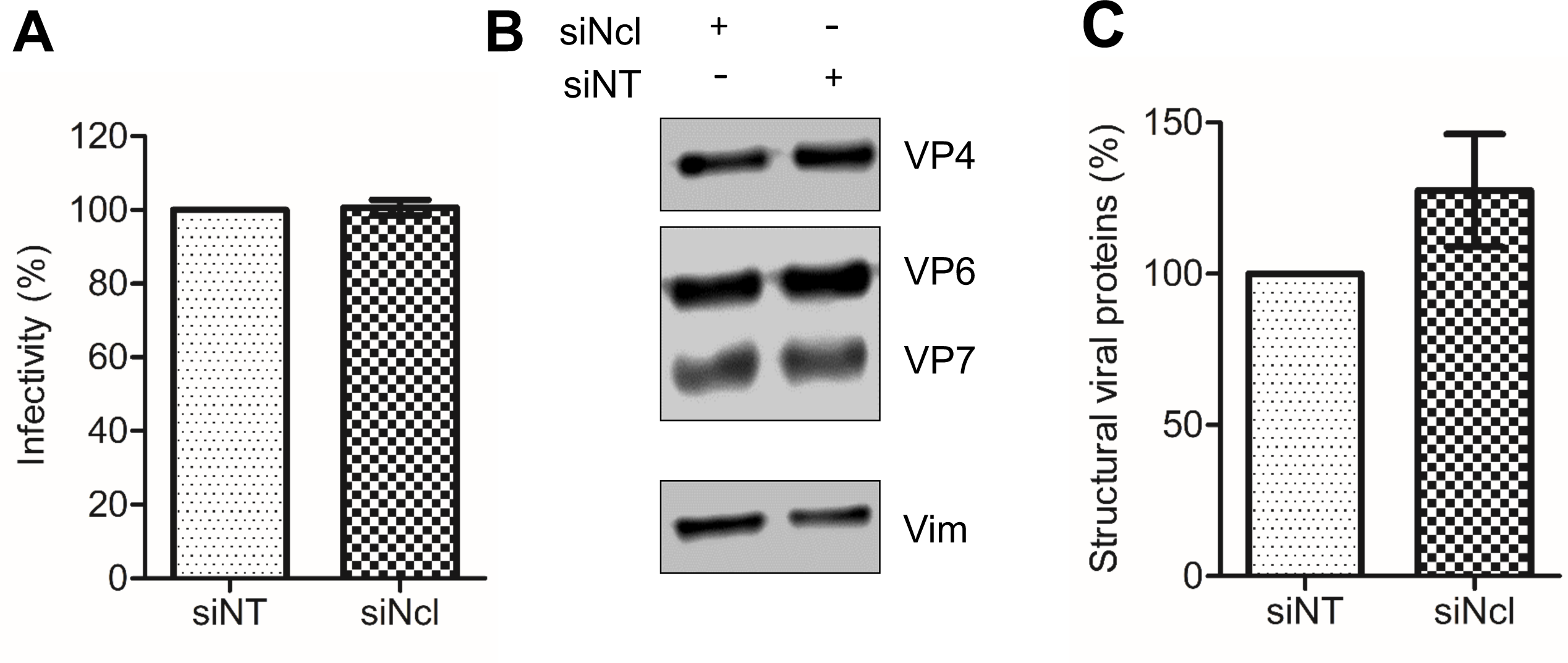
Low nucleolin expression does not have an effect on rotavirus infectivity or viral protein synthesis. A) MA104 cells grown in 96 wells were transfected with either NT or Ncl siRNA as described previously. Cells were then infected with RRV at an MOI of 0.02, and the virus titer was determined at 15 hpi by an immunoperoxidase focus-forming assay, as described under Materials and Methods. The number of infected cells that were transfected with the NT siRNA was taken as 100% infectivity. B) MA104 cells were transfected with either NT or Ncl siRNAs, followed by infection with RRV at an MOI of 3. At 15 hours post-infection cells were harvested, lysed and the proteins were resolved by SDS-10% PAGE, transferred to a nitrocellulose membrane, and detected by immunoblot analysis, using a polyclonal antibody that recognizes rotavirus structural proteins, and an antibody to Vimentin (Vim), as a loading control. C) Densitometry analysis of three rotavirus structural proteins, VP4, VP6, and VP7, and Vim obtained from the Western blot shown in B) was performed using ImageJ software. The arithmetic means ± standard deviation from three independent experiments.

To assess the impact of nucleolin on the synthesis of viral proteins we analyzed the amount of viral proteins by characterizing the abundance of three structural viral proteins (VP4, VP6, and VP7) by densitometry, in lysates of cells in which nucleolin was knocked down. MA104 cells were transfected with nucleolin siRNA, and subsequently infected at 72 hpt with rotavirus RRV at an MOI 3. Cells were harvested at 15 hpi, and the infected cell lysates were analyzed by Western blot SDS-PAGE using an anti-TLP polyclonal antibody. No statistically significant difference was observed between the control NT-siRNA treated cells and the nucleolin-silenced cells (Fig. 2C). These results confirm that the amount of rotavirus proteins remains unaffected in the context of reduced expression of nucleolin.

### Rotavirus genome replication is enhanced in cells in which the expression of nucleolin was silenced

The synthesis of negative-sense RNA during rotavirus infection is a measure of genome replication since this is a mandatory step for the synthesis of the genomic double-stranded RNA present in the infectious viral progeny. To quantify the synthesis of genomic RNA in RRV-infected cells that had been previously treated with nucleolin siRNAs, the specific abundance of the negative strand of rotavirus segment 10 (S10) was amplified by RT-qPCR as an indicator. For this, MA104 cells knocked down for nucleolin were infected with RRV at an MOI of 3 and total RNA was extracted from samples collected at 0, 4, 8, and 12 hpi. Data is expressed as fold increases in which 12 hpi as one in control cells that were transfected with the NT siRNA, since previous observations in our laboratory have indicated that the maximum amount of viral RNA is produced at that time (Ayala-Bretón et al., 2009). To quantitate the amount of S10 negative and positive strands we used a standard curve for the qPCR analysis, using a plasmid construct containing the cDNA of RRV S10 gene segment (see Material and Methods). The synthesis of viral negative-strand under conditions where the expression of nucleolin was silenced showed an increase starting at 4 hpi, remaining sustained until 12 hpi, and showing a substantial 5.8-fold increase compared with the RNA synthesis observed in control cells treated with the NT-siRNA (Fig. 3A).

**Figure 3.**
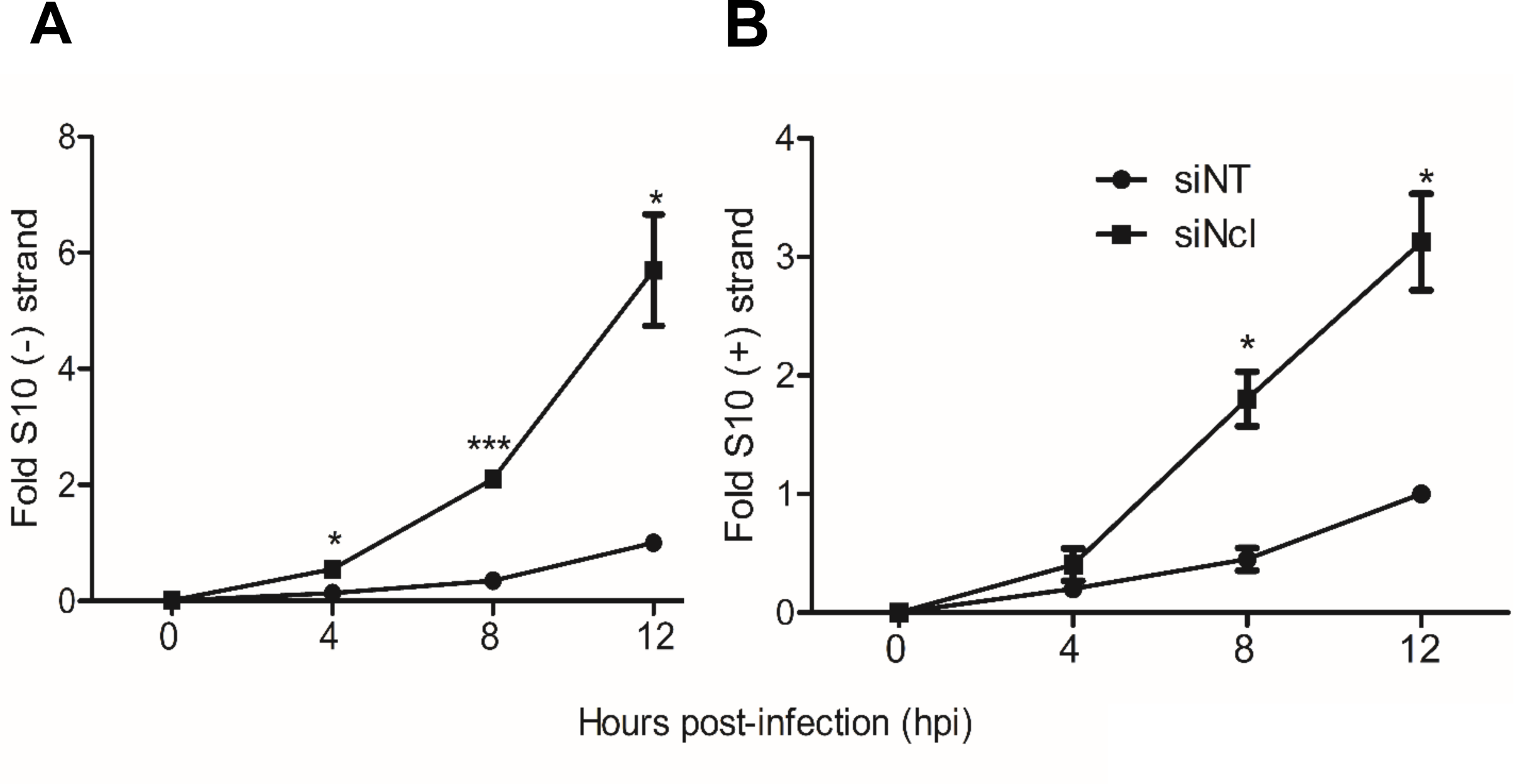
The synthesis of the negative and positive strands of viral RNA are enhanced in Nucleolin-Silenced Cells. A) MA104 cells grown in 48 wells were transfected either with NT or Ncl siRNA as previously described, and subsequently infected with RRV at an MOI of 3. Samples were collected at the indicated hpi, and total RNA was extracted using Trizol. The levels of both strands of the RRV S10 were determined by real-time RT-PCR as described under Materials and Methods. Data is expressed as fold increases in which the RNA copies obtained from the cells transfected with the NT siRNA and infected at 12 hpi was used to normalize. A) S10 negative strand levels and B) S10 positive strand levels. The arithmetic means ± standard deviation of three independent experiments is presented.

We also determined the abundance of the S10 positive strand, which represents the sum of the double-stranded RNA genome synthesis and the mRNA synthesized from the transcription process. Similar to what we observed with the S10 negative strand, an increase in the abundance of the S10 RNA positive strand was observed at 8 hpi compared to control silenced cells, with a significant 3.5-fold enhancement at 12 hpi (Figure 3B). These findings support the notion that under conditions in which nucleolin expression is silenced, there is an increased viral RNA synthesis that contributes to the observed increase in viral progeny production.

### Nucleolin interacts with rotavirus RNA segment 10 during infection

To demonstrate the potential interaction between nucleolin and rotavirus RNA, an immunoprecipitation assay (IPP) was conducted. In this assay, a monoclonal antibody against nucleolin was utilized to immunoprecipitate infected or mock-infected cell lysates. The presence of rotavirus RNA in the immunoprecipitated complex was then detected using RT-PCR to amplify the rotavirus RNA S10 segment. As positive control, an immunoprecipitation using an antibody to the virus nonstructural protein NSP3 was included, since it is known that this protein specifically binds to all rotavirus RNA segments (López & Arias, 2012). Additionally, as a negative control, an antibody to Protein phosphatase 1 (PP1), that has been previously shown that it does not interact with rotavirus RNA (Oceguera et al., 2018), was used.

The effectiveness of the IPP assays was verified by Western blot analysis of the precipitates using antibodies specific for nucleolin, NSP3, and PP1 (Figure 4A). When the nucleolin monoclonal antibody was used to IPP putative protein-RNA complexes in cell lysates, an RT-PCR amplicon of 751 bp, corresponding to the S10 segment of rotavirus, was obtained (Figure 4B) from infected cells. This amplicon was also observed when the polyclonal antibody to NSP3 was used in the IPP. Importantly, no amplification bands were observed when using the PP1 antibody. Altogether these results provide evidence for a specific interaction between nucleolin and the RRV S10 mRNA.

**Figure 4.**
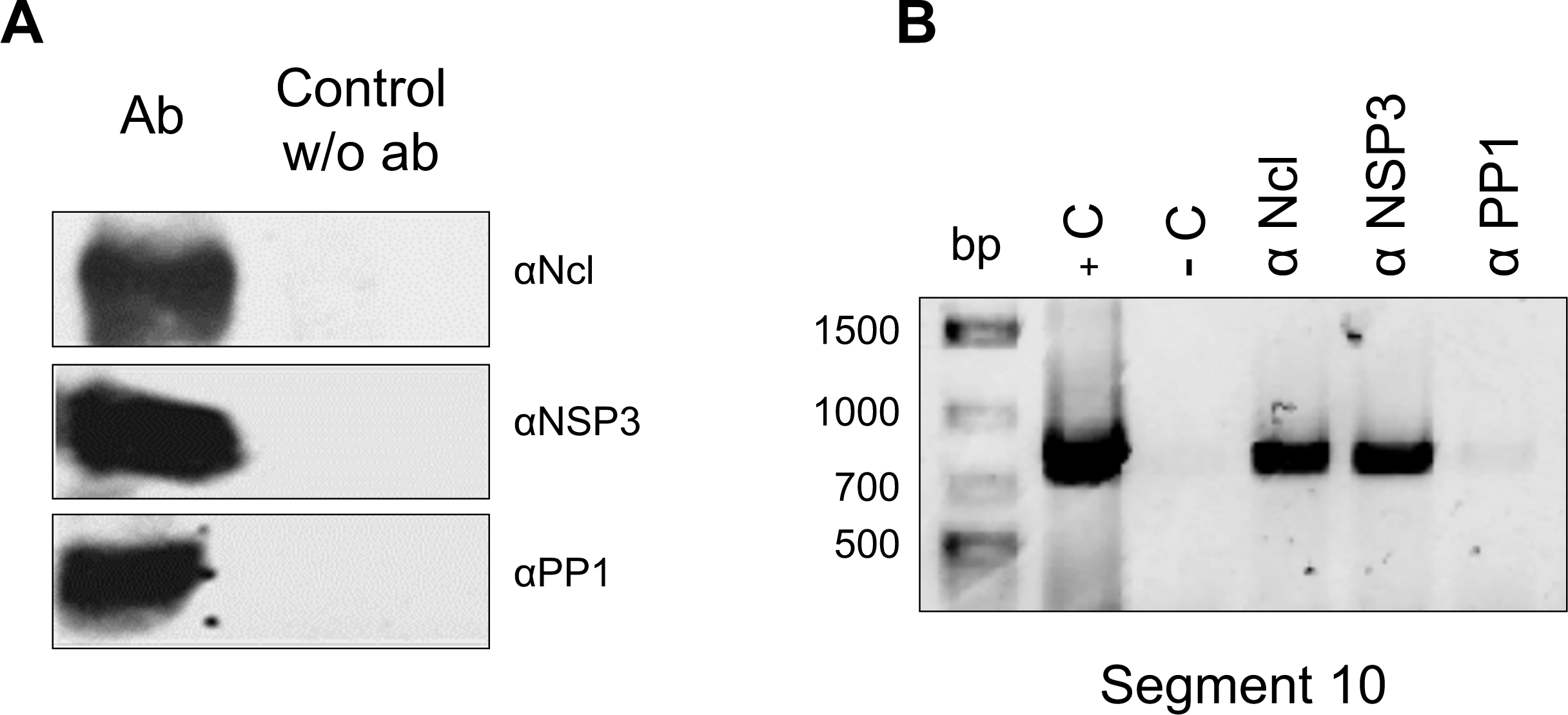
Nucleolin binds to viral RNA in rotavirus infected cells. Confluent MA104 cells grown in 6 well plates were infected at an MOI of 5 and harvested at 8 hpi. Cells were lysed by ultrasonication, and the cell lysates were used for immunoprecipitation (IPP) using the indicated monoclonal or polyclonal antibodies, or mock IPP without added antibody (w/o Ab). A) The immunoprecipitated were resolved in an SDS-10% PAGE and transferred to nitrocellulose membrane. The proteins were detected by immunoblot analysis with the indicated antibodies. B) The RNA present in the IPPs was extracted with Trizol and RRV segment 10 was amplified by RT-PCR. The amplicon was resolved in 1% agarose gel. As template, RNA from rotavirus was used as positive control (+C) and water as negative control (-C) of reaction.

### The presence of parallel G4 structures in rotavirus RRV genomic segments is identified *in silico*

The potential interaction between nucleolin and viral S10 RNA may be attributed to the presence of RNA-binding domains present in nucleolin. However, the precise regions of viral RNA that could potentially engage in these interactions have yet to be elucidated. Previous studies have highlighted that G-quadruplex domains (G4) present in the RNAs of various viruses can interact with nucleolin (Tosoni et al., 2015; Lista et al., 2017; Bian et al., 2019).

To determine the presence of putative G4 regions within the rotavirus genome, we employed the bioinformatic tool QGRS mapper, as previously described (Dabral et al., 2019). We analyzed all eleven rotavirus RRV mRNAs. Table 1 shows the summary of the results obtained, which reveal the presence of G4 regions in every genomic RNA segment. Closer analysis of rotavirus S10 identified three potential G-4 binding sequences for nucleolin (designated as G4-1, G4-2, and G4-3). G4-1 (nt 116-132) was found near the 5’ UTR region, G4-2 (nt 572-607) was located in the open reding frame (ORF), and G4-3 (nt 717-737) was situated in the 3’ UTR region.

**Table 1.**
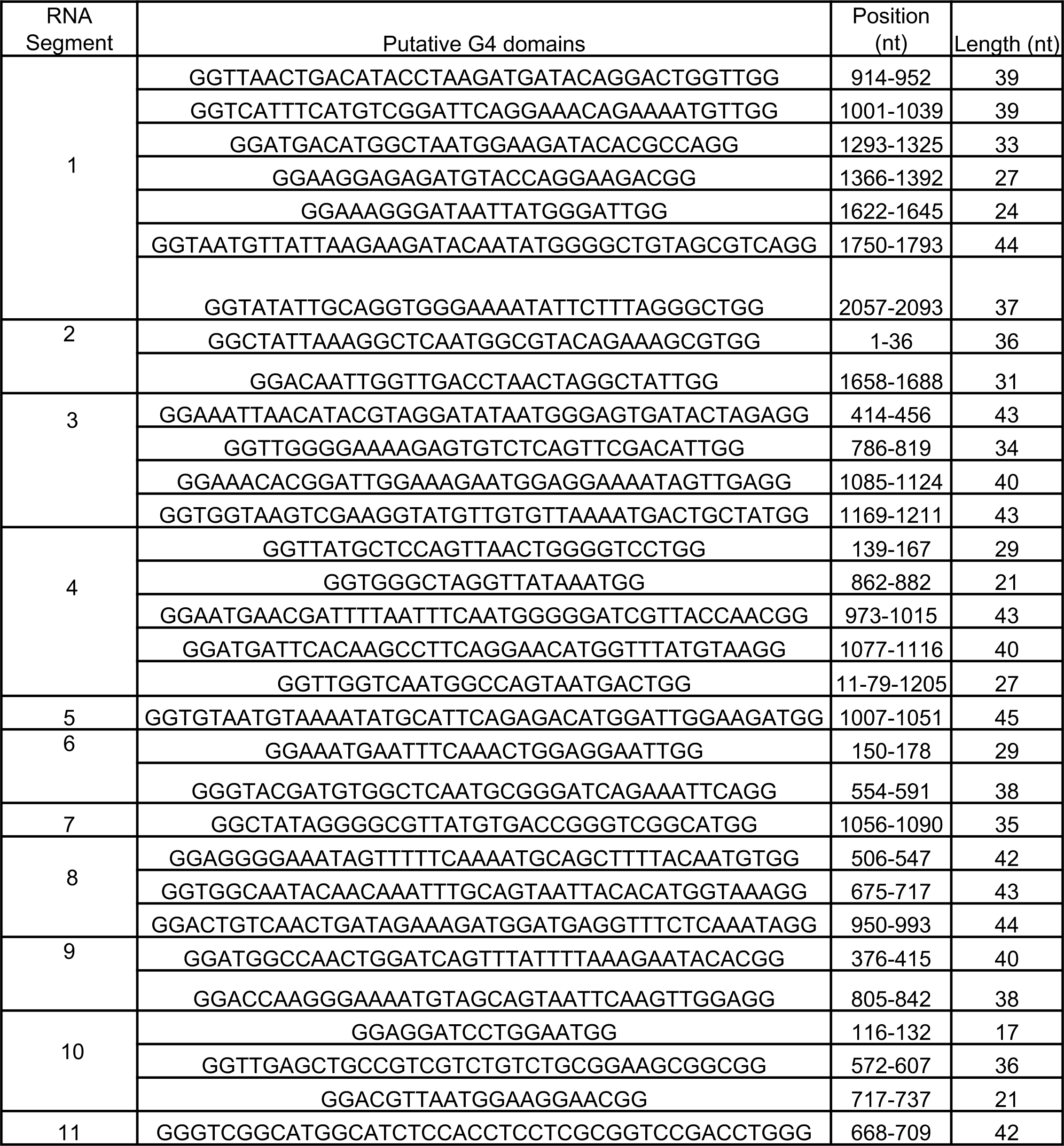
Sequences of predicted G4 sequences in the eleven RNA genome segments of RRV using QGRS mapper web server. The sequence of the positive polarity of all RNA segments is shown.

### Nucleolin restricts RRV synthesis of infectious viral progeny by interacting with G-quadruplex structures present in the viral RNAs

To determine if nucleolin binding to rotavirus S10 RNA is mediated by interactions with the G4 domains identified, MA104 cells were transfected with AGRO100, an oligonucleotide known to form G-quadruplex structures and to specifically bind to nucleolin’s RNA binding domains (Girvan et. al; 2006). As a negative control, cells were transfected with CRO26, an oligonucleotide with the same sequence of AGRO100, in which the guanines were replaced by cytosines. In these assays, MA104 cells previously transfected with either oligonucleotide were infected with RRV at an MOI of 3, and at 15 hpi the infected cells were collected, and the production of viral progeny was determined as previously described. We found that there was a 1.5-fold increase in viral production in the cells transfected with AGRO100 as compared to the control cells transfected with CRO26 (Figure 5A). This finding suggests the involvement of nucleolin’s RNA binding motif in the regulation of rotavirus infectious particle production.

**Figure 5.**
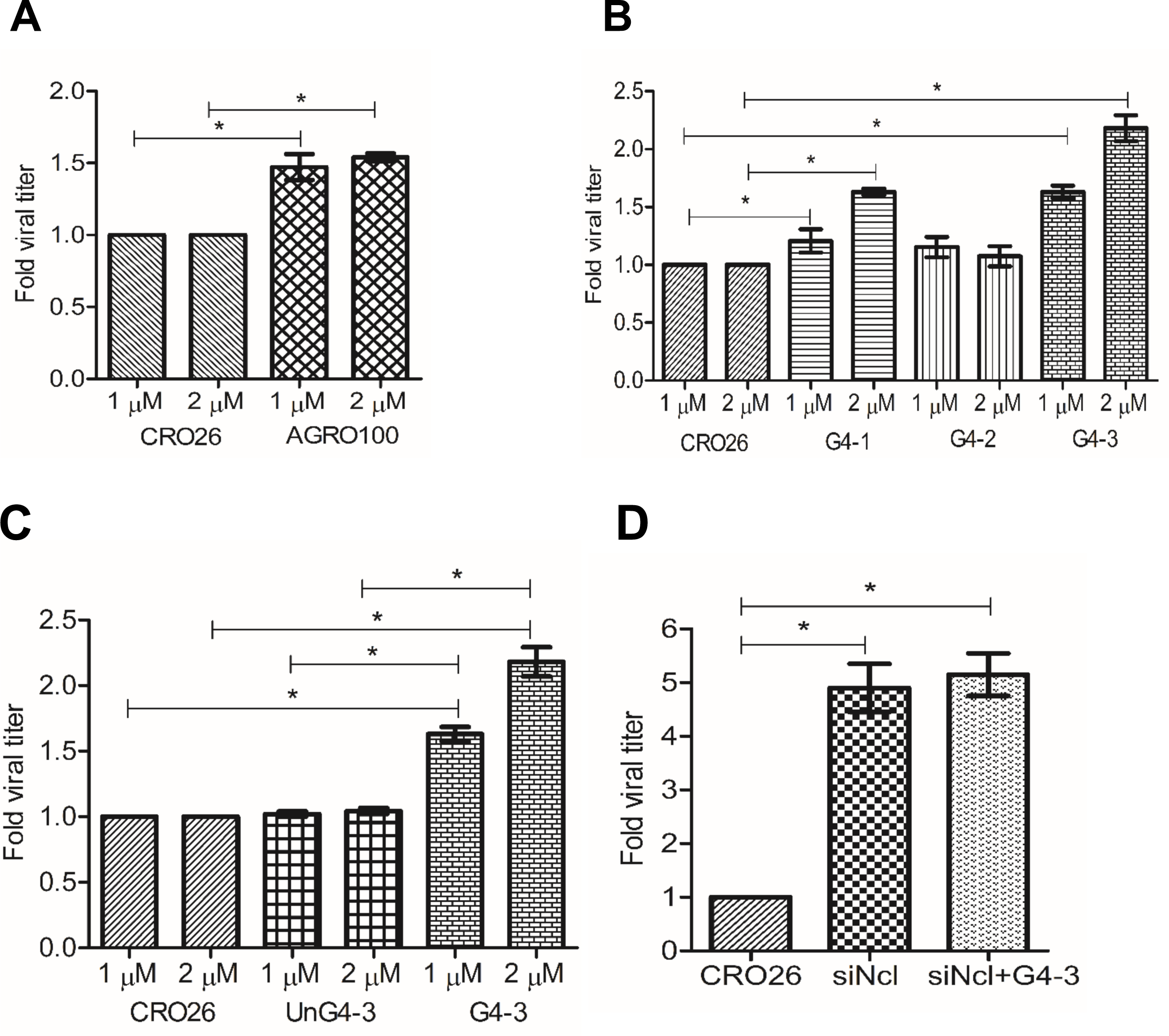
Effect of oligonucleotides with predicted G4 structures on the production of rotavirus progeny. MA104 cells grown in 48 wells were transfected with different oligonucleotides sequence, using the indicated concentrations, for 48 hpt. Transfected cells were infected with RRV at a MOI of 3. After 15 hpi, the virus produce was harvest and the viral titers were determined by an immunoperoxidase foci formation assay as described under Materials and Methods. A) CRO26 and AGRO100. B) Sequences of the three different putative G4 regions present S10 RRV: G4-1 (116-132 nt), G4-2 (572-607 nt), and G4-3 (717-737 nt). C) CRO26, G4-3 unfolded sequence (UnG4-3) and G4-3 folded. D) MA104 cells grown in 48 wells were transfected with nucleolin siRNA. At 24 hours post-transfection (hpt), cells Ncl+G4-3 were now transfected with G4-3 sequence at 2 μM. At 48 hpt, all the cells transfected were infected with RRV at a MOI of 3. After 15 hpi, the virus produced was harvest and the viral titers were determined by an immunoperoxidase foci formation assay as described under Materials and Methods. Data is expressed as fold increases in which the titer obtained from the cells transfected with the CRO26 was used as normalizer. The arithmetic means ± standard deviation three independent experiments are presented.

To determine whether the predicted G-quadruplex structures identified in the rotavirus genome could mimic the effects of AGRO100, we synthesized the three G4 sequences identified in RNA S10 (G4-1, G4-2, and G4-3; see Table S1). These sequences, subjected to an *in vitro* folding treatment as detailed under Materials and Methods, were individually transfected into MA104 cells to assess their impact on virus yield. Notably, RRV-infected cells treated with oligos G4-1 and G4-3 showed a significant increase in viral production of 1.6-fold and 2.1-fold, respectively, compared with the control oligonucleotide (CRO26). In contrast, oligo G4-2 did not significantly affect viral production compared to the controls (Figure 5B). As an additional control, oligonucleotide G4-3 was transfected prior to the *in vitro* folding treatment, (UnG4-3). The results of this assay showed no change in the viral titer compared with the negative control CRO26 demonstrating that the folding of the G4 structure is necessary to affect the replication of the virus (Figure 5C).

Finally, to eliminate the possibility that the observed increased infectious virus production could be a consequence of nonspecific interactions of the RRV G-quadruplex regions with other cellular proteins, and to establish a direct association between the increased virus production and the binding of the RRV-G4 sequences with nucleolin, the expression of this protein was silenced by RNAi in MA104 cells, and 24 h post-transfection, cells were subsequently transfected with the folded G4-3 oligonucleotide for 48 h. After this time, the cells were infected with RRV and 15 hpi at 37 °C. Quantification of infectious viral progeny revealed that the virus titer obtained in nucleolin-silenced cells exhibited no statistically significant difference with the cells that were additionally transfected with RRV G4-3 (Figure 5D).

These results show that G-quadruplex structures favor rotavirus RNA replication by blocking nucleolin’s RNA binding domain. Furthermore, they suggest that the specific G-quadruplex sequences found in S10 of RRV (G4-1 and G4-3) might represent binding sites for nucleolin and this interaction could represent one of the antiviral responses of the cell.

## Discussion

Cellular RNA-binding proteins have been extensively studied in the context of viral infection, for viruses with both DNA and RNA genomes (Lisy et. al., 2021). They have been shown to play diverse roles, including the recognition of viral RNA (vRNA) templates, recruitment of vRNA to replication factories, and the switch from translation to replication (Z. Li & Nagy, 2011; Nagy & Pogany, 2011; R. Y. Wang & Li, 2012). Additionally, they have been involved in the nucleus-cytoplasm shuttling, in particular, hnRNPs such as PCBP1, PTBP1, HNRNPA1, HNRNPC, and nucleolin have been extensively studied (Lloyd et al., 2015). It has been shown that nucleolin plays a significant role in various stages of the replication cycle of several RNA genome viruses (Balinsky et al., 2013; Bian et al., 2019; Bose et al., 2004; Cancio-Lonches et al., 2011; Hernández et al., 2016).

In the specific case of rotaviruses, several studies have reported interactions between cellular RNA binding proteins and vRNA, for example has been shown that heterogeneous nuclear ribonucleoproteins (hnRNPs) D, E, H, and L are relocated from the nucleus to the cytoplasm through interactions with the viral proteins NSP2 and NSP5; furthermore, knockdown of hnRNPs D, I, and L cause a decrease in viral progeny production. In contrast, knockdown of hnRNP C1 and E, produce an increase in viral progeny (Dhillon et al., 2018). Our laboratory investigations have demonstrated that during rotavirus infection the human antigen R protein (HuR) is relocalized to the cytoplasm, while GW182 and argonaute2 (Ago2) proteins accumulate around viroplasms; the relocalization effect of these cellular proteins was proposed to be due to a sponge effect of the viral RNA of rotavirus (Oceguera et al., 2018). Furthermore, ATP synthase has been identified to interact with the 3’ untranslated region (UTR) of the rotavirus genome and to co-localize with viral RNA and viroplasms. Remarkably, knocking down the expression of ATP synthase F1 subunit beta (ATP5B), using RNA interference (RNAi) led to a reduction in the production of infectious viral progeny (Ren et al., 2019).

In this study, we show that silencing the expression of nucleolin in MA104 cells results in an increase in viral mRNA synthesis and genome copy number. The knockdown of nucleolin also resulted in a 5-fold increase in the number of infectious viral particles produced, suggesting that nucleolin down-regulates several processes like replication and viral particle production. Importantly, the decrease in nucleolin expression did not affect the number of infected cells, indicating that the enhanced viral titer production is independent of changes in the cells’ susceptibility to the infection.

Interestingly, no statistically significant change in the synthesis of rotavirus proteins was observed, consistent with previous findings from our laboratory in which it was found that knocking down the expression of NSP3 resulted in an increased viral mRNA and dsRNA levels, without significantly impacting the level of viral protein synthesis (Montero et al., 2006). Furthermore, silencing the expression of VP1 and VP3 resulted in a decrease of about 90% in the levels of viral mRNA, yet the synthesis of viral proteins was not affected (Ayala-Breton et al., 2009). Taken together these results support the notion that the amount of viral mRNA present in the infected cell and the translation of viral proteins are not necessarily directly proportional.

A similar enhancement on the synthesis of viral RNA induced by knockdown of nucleolin expression has been previously reported for hepatitis C (HCV) (Bian et al., 2019) and dengue (Phillips et al., 2016) viruses; in the latter case, an increased production of infectious virus particles was also observed. In addition, the interaction of nucleolin with viral RNA has been reported for human immunodeficiency virus (HIV) and Epstein-Barr virus. In the case of HIV, it was found that nucleolin binding to the virus LTR through a G4 sequence led to an increased LTR promoter activity (Tosoni et al., 2015). For Epstein-Barr virus, nucleolin binds to the EBNA1 mRNA, repressing its translation (Lista et al., 2017). Finally, nucleolin binds to core gene of HCV through a G4 domain and inhibited RNA replication (Bian et. al.; 2019).

In this work, using the QGRS mapper we found that all eleven rotavirus RNA segments contain putative G4 domains suggesting a potential mechanism of interaction between nucleolin and rotavirus RNA segments. This interaction was experimentally confirmed by co-ipp of nucleolin and rotavirus RNA S10. It has been shown that nucleolin specifically recognizes G4 structures in the RNAs it binds to. In this regard, we showed that AGRO100, an oligonucleotide that forms a G4 structure increases viral yield when transfected into cells before virus infection, most probably by binding to nucleolin and preventing it from interacting with viral RNA. Furthermore, and most interestingly, when oligonucleotides representing the predicted G4 sequences in rotavirus genomic S10 were transfected into cells before RRV infection, an increase in viral progeny production, similar to that induced by knocking down the expression of nucleolin, was observed. Altogether these findings suggest that nucleolin down-regulates the synthesis of rotavirus RNAs, with the consequent reduction of infectious viral yield, by binding to G4 structures, probably on RNAs derived from all eleven virus segments.

Of interest one of the three oligonucleotides containing S10 G4 sequences (G4-2), as opposed to the other two (G4-1 and G4-3) did not promote the increase in virus replication. This discrepancy could be attributed to the larger size of the G4-2 sequence and its impossibility to form G-quadruplex, preventing its recognition by the RNA-binding domains of nucleolin.

The observation that there is no statistical difference between the production of viral particles in cells in which nucleolin expression was silenced and transfected with rotavirus G4 sequences, compared to cells where only nucleolin was silenced, suggests that the impact of the G-quadruplex regions present in rotavirus RNA is specific to block the RBD of nucleolin. If it were an unspecific effect, we would have expected to see a combined increase in viral particle production in the double transfection, nucleolin silencing and G-quadruplex transfection. However, no significant difference was observed between these two experimental conditions.

Proteins with RNA-binding domains, including nucleolin, can also exhibit antiviral effects by binding to exogenous RNA and triggering immune responses. For instance, ZNFX1 can bind to viral double-stranded RNA (dsRNA) and interact with MAVS, leading to the promotion of interferon expression (Wang et al., 2019). Although it remains to be explored, it would be interesting to investigate whether nucleolin plays a similar role in the antiviral response. Additional experiments such as co-immunoprecipitation or protein-protein interaction assays should be performed to assess the presence of any potential proteins involved in the nucleolin-rotavirus RNA interaction.

In previous reports from our laboratory, it was found that during rotavirus infection there is a substantial accumulation of viral mRNA [for example, at 8 hpi it was estimated that the NSP4 mRNA reached 1 × 10^5 copies per cell (Ayala-Breton et al., 2009)], and it has been proposed that these amount of viral RNA could be sequestering many of the RBP of the cell (Oceguera et al., 2018). The concept of an “RNA sponge” effect, derived from this observation, may provide insights into how rotavirus circumvents the inhibitory influence of nucleolin described in this work, thereby allowing the presence of vRNA free from nucleolin interactions.

In conclusion, this study emphasizes the importance of nucleolin in rotavirus infection by negatively regulating viral RNA synthesis and infectious virus production. Further research is warranted to deepen our understanding of the intricate interactions between nucleolin and rotavirus during infection. The field of research on proteins with RNA-binding domains is expected to expand due to their relevance in RNA virus infection.

## Acknowledgements

This work was supported by the Universidad Nacional Autónoma de México. We thank P. Gaytán Colín, E. López-Bustos, and S. Becerra Ramírez from the Unidad de Síntesis y Secuenciación de DNA, Instituto de Biotecnología, UNAM, for synthesis of the oligonucleotides used in this work. We are grateful to Rafaela Espinosa, and Marco Antonio Espinoza for their technical assistance.

